# Expression of *Bacillus cereus* Virulence-Related Genes in an Ocular Infection-Related Environment

**DOI:** 10.1101/2020.03.17.995753

**Authors:** Phillip S. Coburn, Frederick C. Miller, Morgan A. Enty, Craig Land, Austin L. LaGrow, Md Huzzatul Mursalin, Michelle C. Callegan

**Author notes:** Corresponding author: Phillip S. Coburn,. Abbreviations: LB, Luria Bertani; BHI, Brain Heart Infusion; RNA-Seq, RNA Sequencing; POE, post-operative endophthalmitis; PTE, post-traumatic endophthalmitis; EE, endogenous endophthalmitis; PC-PLC, phosphatidylcholine-specific phospholipase C; PI-PLC, phosphatidylinositol-specific phospholipase C; TLR, Toll-like receptor; Hbl, hemolysin BL; Nhe, nonhemolytic enterotoxin; Ent, enterotoxin; InhA, immune inhibitor A; CerO, cereolysin O; HylA, hemolysin A; SOD, superoxide dismutase; RPKM, reads per kilobase million, CT, threshold cycle; bp, basepairs; ANOVA, analysis of variance.

## Abstract

*Bacillus cereus* produces many factors linked to pathogenesis and is recognized for causing gastrointestinal toxemia and infections. *B. cereus* also causes a fulminant and often blinding intraocular infection called endophthalmitis. We reported that the PlcR/PapR system regulates intraocular virulence, but the specific factors that contribute to *B. cereus* virulence in the eye remain elusive. Here, we compared gene expression in *ex vivo* vitreous humor with expression in Luria Bertani (LB) and Brain Heart Infusion (BHI) broth by RNA-Seq. The expression of several cytolytic toxins in vitreous was less than or similar to levels observed in BHI or LB. Regulators of virulence genes, including PlcR/PapR, were expressed in vitreous. PlcR/PapR was expressed at low levels, though we had reported that PlcR-deficient *B. cereus* was attenuated in the eye. Chemotaxis and motility genes were expressed at similar levels in LB and BHI, but at low to undetectable levels in vitreous, although motility is an important phenotype for *B. cereus* in the eye. Superoxide dismutase, a potential inhibitor of neutrophil activity in the eye during infection, was the most highly expressed gene in vitreous. Genes previously reported to be important to intraocular virulence were expressed at low levels in vitreous under these conditions, possibly because *in vivo* cues are required for higher level expression. Genes expressed in vitreous may contribute to the unique virulence of *B. cereus* endophthalmitis, and future analysis of the *B. cereus* virulome in the eye will identify those expressed *in vivo*, which could potentially be targeted to arrest virulence.

**Impact statement:** *B. cereus* is the causative agent of gastrointestinal infections, but can also cause a serious infection of the eye that often results in blindness or enucleation. Current therapeutic measures often fail to mitigate these poor outcomes. This necessitates the development of new treatment modalities based on new targets. To begin to better define those *B. cereus* factors with roles in intraocular infection, we analyzed the expression of genes related to gastrointestinal infections, as well as those with both known and hypothesized roles in intraocular infections, after growth in an *ex vivo* vitreous. Potentially targetable candidate genes were demonstrated to be expressed in vitreous, which suggests that these genes might contribute to the unique virulence of *B. cereus* endophthalmitis. Importantly, our results lay the groundwork for assessing the expression of these genes *in vivo* and defining the virulome of *B. cereus* in intraocular infections.

## Introduction

*B. cereus* is one of the leading causes of bacterial gastrointestinal infections and produces a variety of toxins that contribute to the pathogenesis of these infections. *B. cereus* also poses a serious threat to vision should it gain access to the interior of the eye. Endophthalmitis is an infection of the anterior and posterior segments of the eye resulting from contamination with microorganisms following a surgical procedure (post-operative endophthalmitis [POE]), a traumatic penetrating injury (post-traumatic endophthalmitis [PTE]), or metastasis from an infection of a distant site in the body (endogenous endophthalmitis [EE]) [1–5]. *B. cereus* is a leading cause of both PTE and EE. These infections result in a fulminant endophthalmitis characterized by severe intraocular inflammation, ocular pain and proptosis, and significant vision loss within hours [1–5]. The significant ocular damage occurring during *B. cereus* endophthalmitis is presumably due to a combination of bacterial-mediated and host immune-related mechanisms. The majority of patients afflicted with this disease (∼70%) lose significant vision, if not the eye itself, in a few days, regardless of treatment measures [1–5]. *B. cereus* endophthalmitis is often refractory to treatment because of the rapid nature of the infection. As a whole, endophthalmitis can be difficult to treat due to ineffective antibiotic penetration, infection with antibiotic-resistant organisms, conflicting clinical information with regards to the dose, route, and combination therapy, or delays in time between injury and treatment. Therapies aimed at preserving visual acuity are often inadequate for *B. cereus* endophthalmitis and, at best, can prevent enucleation of the eye. A better understanding of the mechanisms and factors involved in the pathogenesis of *B. cereus* PTE and EE is therefore urgently needed.

During endophthalmitis, *B. cereus* toxins may injure the nonregenerative tissues of the eye directly by actively damaging cells or indirectly by inciting inflammation that damages or interferes with the physiological processes of vision [6–10]. When groups of toxins are absent, such as those regulated by the PlcR/PapR transcriptional regulatory system, the virulence of endophthalmitis is significantly muted [9]. However, our previous analyses of the contributions of hemolysin BL, phosphatidylcholine-specific phospholipase C (PC-PLC), and phosphatidylinositol-specific phospholipase C (PI-PLC) did not reveal individual roles for these toxins in a rabbit model of endophthalmitis [7, 11]. The specific toxins of *B. cereus* which contribute to its unique virulence in the eye remain an open question.

Rapidly evolving endophthalmitis caused by *B. cereus* has also been attributed to cell wall and envelope associated factors that activate a greater inflammatory response than do other intraocular pathogens [12]. The cell envelope of *B. cereus* consists of a thick peptidoglycan layer associated with a capsular polysaccharide, lipotechoic and teichoic acids, lipoproteins, pili, a glycoprotein S-layer, and flagella [13–15]. The cell wall of *B. cereus* incited a greater inflammatory response than the cell walls of the intraocular pathogens *S. aureus* and *E. faecalis* following injection into a rabbit eye [12]. This suggests that either *B. cereus* possesses a unique cell wall component(s) that strongly activate(s) an inflammatory response, or structural differences of a shared component increase the ability of *B. cereus* to incite a response. The presence of pili appeared to protect *B. cereus* from clearance in the mouse eye [16], suggesting that pili might function as an antiphagocytic factor in the intraocular environment. A unique structural feature that distinguishes the *B. cereus* cell wall from the cell walls of the other leading Gram-positive causes of endophthalmitis is the S-layer. We recently reported its significant contribution to the intraocular inflammatory response [17]. Flagella aid *B. cereus* in migration through the eye [12, 18], but the *B. cereus* flagella does not activate Toll-like receptor (TLR)-5, the innate immune receptor which recognizes flagella [19]. However, we demonstrated that flagellar motility and the swarming phenotype are important to the intraocular virulence of *B. cereus* [8, 18]. *B. cereus* rapidly migrates throughout all parts of the eye, and elicits significant and damaging inflammation [12]. However, attenuation by mutating motility phenotypes only delayed the evolution of disease [8, 18], suggesting that other factors are involved in virulence. The devastating nature of *B. cereus* endophthalmitis [1–5] warrants the identification of better targets for treatment of the disease.

The robust inflammatory processes which occur in response to *B. cereus* infection are triggered by the initial recognition of cell wall/envelope components and secreted products via a class of pattern recognition receptors called Toll-like receptors (TLRs) that are expressed on host cells [20, 21]. During the early stage of *B. cereus* endophthalmitis in mice, a number of retinal genes were upregulated, including many associated with the host acute inflammatory response. Fifteen genes upregulated 5-fold or higher following infection were associated with TLR4-induced inflammation [22], demonstrating that *B. cereus* is capable of activating TLR4 in a mouse model of endophthalmitis. The explosive nature of *B. cereus-*induced endophthalmitis suggests that virulence-related *B.* cereus genes are expressed in the vitreous environment and induce a rapid and destructive immune response.

In the current study, we evaluated the expression levels of a subset of factors involved in virulence and its transcriptional regulation, motility, and chemotaxis after growth to stationary phase in an ocular infection-related environment *ex vivo.* This late stage correlates with a stage of infection when significant inflammation and retinal function loss have occurred. Characterizing the expression of this subset of virulence-related genes in an ocular infection-related environment might provide insight into the factors that are responsible for the devastation to the eye during the later stages of infection. This was accomplished by analyzing the expression levels of virulence-related genes of *B. cereus* grown to stationary phase in explanted vitreous compared to standard laboratory media using RNA-Seq. In all environments tested, we observed a continuous spectrum of gene expression from low to high levels. Across all genes surveyed, normalized expression levels tended to be lower in vitreous relative to the laboratory media. Among the genes surveyed, the gene encoding superoxide dismutase, *sodA2*, was the most highly expressed gene in vitreous, whereas genes related to motility and chemotaxis were expressed at the lowest levels. Varying levels of toxin gene expression were also observed in the vitreous during stationary phase. Our results suggested that the *B. cereus* superoxide dismutase might represent a target for therapeutic intervention, and demonstrated that virulence-related genes are expressed in *ex vivo* vitreous during the stationary phase of growth, which may relate to expression in later stages of infection. These results provided the framework for studying the *B. cereus* virulome during ocular infections in order to identify virulence-related genes expressed *in vivo,* and suggest possible candidate genes that could be potentially targeted to mitigate virulence.

## Methods

### Bacterial strain, growth environments, and growth curves

Vegetative *B. cereus* ATCC 14579 was cultivated in Luria Burtani (LB) for 18 h at 37°C. The culture was centrifuged for 10 minutes at 4,300 x g, and the bacterial pellet washed 3 times with sterile phosphate-buffered saline (PBS) pH 7.4 to remove all traces of LB. After the third wash, the bacterial pellet was resuspended in an equal volume of PBS as the original culture volume and diluted to 10^3^ CFU/ml in freshly-prepared brain heart infusion (BHI) broth, LB broth, or *ex vivo* rabbit vitreous and incubated at 37°C for 18 h. Rabbit vitreous was prepared from batches of 50 mature rabbit eyes obtained from Pel-Freez Biologicals (Rogers, AR). An incision was made in the corneal limbus, the aqueous humor and lens discarded, and the vitreous collected from each eye. Vitreous from all 50 eyes was pooled and filter-sterilized with a 0.22 µm Stericup Quick Release Durapore PVDF filtration unit (EMD Millipore Corporation, Billerica, MA). For growth curve analysis, after dilution to 10^3^ CFU/ml in either BHI, LB, or rabbit vitreous, 20 ul aliquots were obtained from each culture and diluted 10-fold in sterile PBS. Aliquots from each dilution were plated onto BHI agar plates for bacterial quantification. Two separate growth analysis experiments were performed using independent batches of BHI, LB, or rabbit vitreous, and each experiment was performed with 3 independent cultures in each environment.

### RNA preparation and quantitative PCR Analysis

For each growth condition, three independent cultures were prepared and RNA was isolated and prepared from each as follows. Cultures were centrifuged for 10 minutes at 4,300 x g and the cell pellet resuspended in the lysis buffer (RLT) from the RNeasy kit (Qiagen, Germantown, MD). Cells were then homogenized with sterile 0.1 mm glass beads (Biospec Products Inc., Bartlesville, OK) for 60 seconds at 5,000 rpm in a Mini-BeadBeater (Biospec Products Inc., Bartlesville, OK). Total RNA was purified using the RNeasy kit according to the manufacturer’s instructrions (Qiagen). Genomic DNA was removed using the TURBO DNA-free kit (ThermoFisher Scientific, Inc., Waltham, Massachusetts). Ribosomal RNA is the most abundant RNA species in the bacterial cell and would interfere with sequencing relatively rare mRNA transcripts. Therefore, ribosomal RNA was depleted using the Ribo-Zero rRNA Removal kit for bacterial rRNA (Illumina, San Diego, CA) according to the manufacturer’s instructions, and depletion was assessed via quantitative PCR using primers specific to the *B. cereus* 16s ribosomal RNA. 100 ng aliquots of RNA before and after depletion were subjected to qPCR using the iTaq™ Universal SYBR^®^ Green One-Step kit (Bio-Rad, Hercules, CA) [22]. The forward and reverse primers were used at a final concentration of 300 nM. The samples were run on a Bio-Rad^®^ CFX96 Touch™ Real-Time PCR System (Bio-Rad) [22]. Dissociation curves were used to assess the successful amplification of the desired product, and the threshold cycle (CT) was used to determine relative amounts of transcripts between RNA samples before and after depletion. RNA fold decreases were calculated by subtracting the CT values of the RNA samples prior to depletion from the CT values of the RNA samples following depletion. That value as a power of 2 yielded the fold decrease of rRNA in the depleted sample relative to the undepleted sample. Reported fold decreases represent the mean fold decrease ± the standard deviation of the three independent RNA samples for each condition.

### RNA sequencing

The triplicate enriched RNA samples obtained from each of the growth conditions was sequenced using an Illumina MiSeq Next Generation Sequencer at the OUHSC Laboratory for Molecular Biology and Cytometry Research. Raw data for each sample was analyzed using the CLC Genomics Workbench software (Qiagen, Redwood City, CA). Raw sequence reads were mapped to the *B. cereus* ATCC 14579 reference genome for identification of the genes of interest in our study expressed under each condition. Reads not mapped were excluded further analysis. The number of reads per gene was normalized according to the total number of reads in each library and the gene size. The resulting number was expressed as normalized reads per kilobase per million (RPKM). The values represent the mean RPKM ± the standard deviation of the three independent sequencing runs for each condition.

### Statistics

For the growth curves, data are the arithmetic means ± the standard deviations of the CFU/ml values from two independent experiments with 3 biological replicates of each condition per experiment. For RNA-Seq, the data are the arithmetic means ± the standard deviations of the RPKM values of each gene derived from three cultures for each condition that were sequenced independently. Comparative differences between groups were taken to be statistically significant when p < 0.05. The one-way ANOVA test followed by Tukey’s post-hoc analysis was used to compare the growth curves and the RPKM values of the genes in each environmental condition. All statistical analyses were performed using GraphPad Prism 8.2.0 (GraphPad Software, Inc., La Jolla CA).

### Data availability

The RNA-Seq data was deposited in the Sequence Read Archive at NCBI. The submission ID # is SUB6908082, the BioProject ID # is PRJNA604224, and the BioSample ID # is SAMN13958353.

## Results

### Growth of *B. cereus* in BHI, LB, and rabbit vitreous

In vitro growth of B. cereus ATCC 14579 in the 3 environments was compared to assess whether differences in growth might impact the results of the RNA-Seq experiments. *B. cereus* ATCC 14579 was cultivated in LB for 18 h at 37°C. The culture was washed 3 times in PBS to remove all residual LB and then subcultured in either BHI, LB, or rabbit vitreous. Bacterial growth was measured every 2 hours for 18 hours. As shown in Fig.1, no statistically significant differences were observed in growth between BHI, LB, or rabbit vitreous, with exception of growth in LB at 6, 8, and 10 hours. The growth yield was statistically significantly lower at each of these time points relative to BHI or vitreous (p<0.05). At 18 hours, there were no significant differences in growth yield between the 3 different environments (p≥0.0732). We did not observe statistically significant variation in growth of *B. cereus* from one batch of vitreous to another. Moreover, as can be observed from Figure 1, *B. cereus* reached stationary phase at 12 hours in all 3 environments and remained at the same concentration at 18 hours. Therefore, length of time in stationary phase did not vary between the 3 environments. These results demonstrate that growth of *B. cereus* was similar in all 3 conditions.

**Figure 1.**
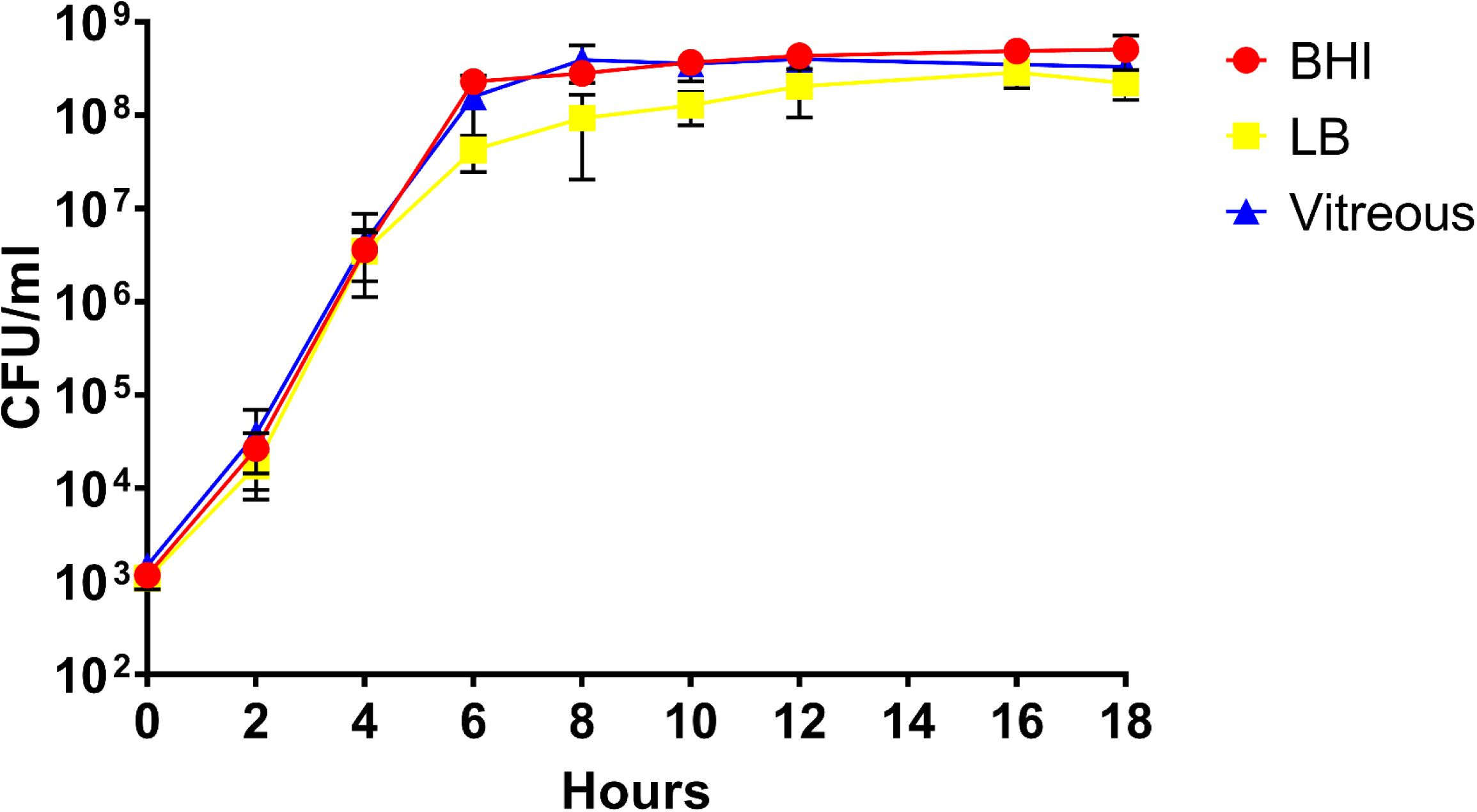
Analysis of the growth of *B. cereus* in BHI, LB, or rabbit vitreous. *In vitro* growth curves of *B. cereus* in the 3 different environments. CFU/ml at each time point was similar in BHI, LB, and rabbit vitreous. The values shown represent the combined results of two separate growth analysis experiments using independent batches of BHI, LB, or rabbit vitreous, with N = 3 independent cultures in each environment per experiment for a total of 6 biological replicates per time point. Values are the mean of the 6 replicates ± SD.

### rRNA depletion and RNA-Seq analysis of depleted RNA samples

To identify genes that might be related to the pathogenesis of *B. cereus* endophthalmitis, we surveyed the expression of a subset of known and putative virulence factors, virulence-associated transcriptional regulators, and genes related to motility and chemotaxis from *B. cereus* after growth in an ocular infection-related environment using RNA-Seq. *B. cereus* ATCC 14579 was studied as a prototype *B. cereus* species with a sequenced and annotated genome. This particular strain has also been well characterized in terms of its virulence and pathogenesis in endophthalmitis [9, 16, 19, 22, 23]. It has 13 rRNA loci, and rRNA constitutes the bulk of the total RNA. As such, rRNA would represent the vast majority of the sequencing reads, and preclude sequencing of relatively rare mRNA species. Therefore, rRNA had to be depleted prior to RNA-Seq analysis. As shown in Table 1, 16s rRNA was depleted by a mean of 5,078-, 8,856-, and 691,802-fold from LB, BHI, and vitreous RNA samples, respectively. These results indicated that rRNA was sufficiently depleted, especially for the vitreous-derived RNA samples.

**Table 1.** Quantitative PCR assessing RiboZero depletion of 16s rRNA in *B. cereus* ATCC 14579.

| Media | RiboZero treated sample<br>(Mean threshold cycle (C<sub>T</sub>)) | Untreated sample<br>(Mean threshold cycle (C<sub>T</sub>)) | rRNA fold depletion |
| --- | --- | --- | --- |
| LB | 19.536 | 7.225 | 5,078 |
| BHI | 17.296 | 4.1825 | 8,856 |
| Vitreous | 25.2 | 5.8 | 691,802 |

The *B. cereus* ATCC 14579 genome possesses 5,473 genes and a length of 5,411,809 bp. The average read length was 132 bp for all samples and conditions. Table 2 shows the mean number of mapped reads and mean percentage of reads that mapped to the *B. cereus* genome for each condition. The mean number of mapped reads for the three BHI-derived RNA samples was 9,609,820 with a mean of 84% of the reads mapping to the *B. cereus* 14579 genome. The mean number of mapped reads for the LB-derived RNA samples was 8,565,599, and a mean of 87% of the reads mapped to the genome. For the vitreous RNA samples, the mean number of mapped reads was 8,350,233 with 94% of the reads mapped. Examination of the *B. cereus* transcriptome revealed that overall, there was a consistent pattern in which gene expression was lower in vitreous relative to LB and BHI.

**Table 2.** Mean number of reads and percentage of reads that mapped to the B. cereus ATCC 14579 genome for each environmental condition.

| Environment | Mean Number of Reads | Percentage of Reads Mapped |
| --- | --- | --- |
| BHI | 9,609,820 | 84% |
| LB | 8,565,599 | 87% |
| Vitreous | 8,350,233 | 94% |

### Virulence factor expression in *ex vivo* vitreous

For the purposes of this study, we focused on known and putative virulence genes, transcriptional regulators of virulence-related genes, and genes related to motility and chemotaxis. We evaluated expression levels of the genes encoding the toxin hemolysin BL, the nonhemolytic enterotoxin Nhe, the putative enterotoxins EntA and EntC, entertoxin FM, the putative hemolysin A, cereolysin O, and the metalloproteases InhA1, InhA2, InhA3, and camelysin. Expression of genes encoding the transcriptional regulatory systems related to virulence (SinR/SinI, EntD, CodY, GntR, NprR, and PlcR/PapR), motility-related proteins Fla, FliF, and MotB, and chemotaxis-associated proteins CheA, CheR, and CheY were also examined. We also sought to determine expression levels of superoxide dismutase in vitreous as this factor might be important to intraocular survival and virulence. Principal component analysis of the RPKM values for each of these mRNAs isolated from the three independent BHI, LB, and vitreous cultures was performed. As can be observed in Fig. 2, with exception of a single LB replicate, the replicates of each condition fell into distinct gene expression clusters, indicating a high degree of reproducibility among replicates.

**Figure 2.**
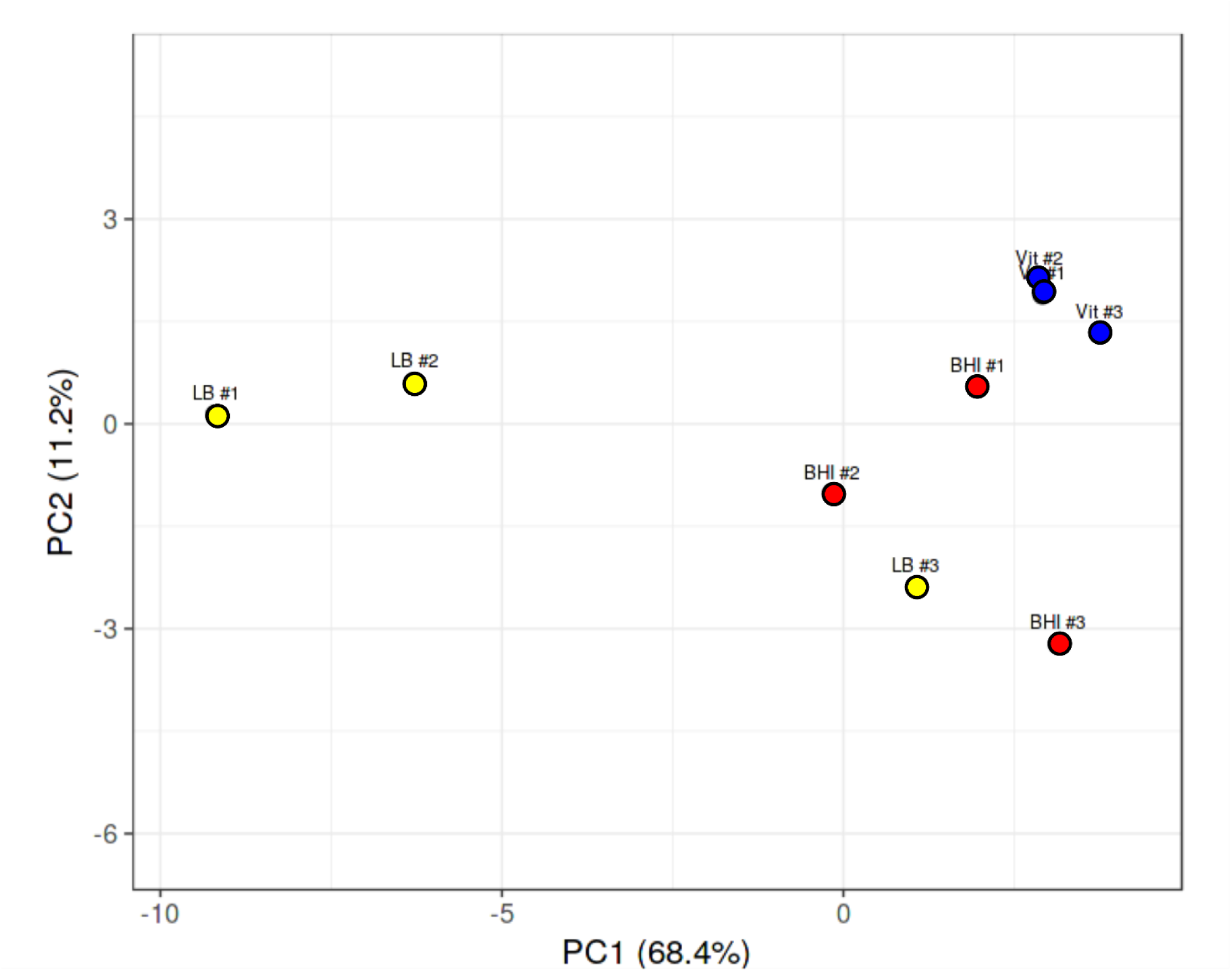
Principal component analysis of *B. cereus* gene expression in LB, BHI, or vitreous. Principal component analysis of expression levels of virulence genes, transcriptional regulators of virulence genes, and genes related to chemotaxis and motility from each of the three independent RNA-seq runs of *B. cereus* cultivated in LB, BHI, or explanted rabbit vitreous at 37°C for 18 h. With exception of one LB run that overlapped with the BHI runs, separate clustering of gene expression according to environment was observed.

Surprisingly, few cytolytic toxins were highly expressed in vitreous. Among those that were expressed were hemolysin BL, the nonhemolytic enterotoxin Nhe, the putative enterotoxins EntA and EntC, entertoxin FM, the putative hemolysin A, cereolysin O, the metalloproteases InhA1, InhA2, InhA3, and camelysin. Mean RPKM values of 62, 65, and 82 were observed for *hblL1*, *hblL2*, and *hblB*, respectively, for the vitreous environment (Fig. 3a). Expression of these genes was 12-, 8-, and 10-fold in higher in BHI (p≤0.0349), and 30-, 30-, and 26-fold higher in LB (p≤0.0264), respectively, relative to vitreous. Similar expression levels of the *nheL1* and *nheL2* genes were observed in the vitreous, with mean RPKM values of 193 and 82, respectively (Fig. 3b). Expression of these genes was not significantly different from BHI (p≥0.9141), but were 8- and 16-fold higher in LB (p<0.05), respectively. Mean RPKM values of 64 for *entA*, and 369 for *entC* were observed in the vitreous (Fig. 3c). Expression of these genes was not significantly different in BHI (p≥0.9823), however were greater in LB by 4-fold for *entA* (p=0.0309), and by 5-fold for *entC* (p=0.0015). Among the enterotoxins, *entFM* was the mostly highly expressed gene in the vitreous with a mean RPKM value of 566 (Fig. 3c). The expression of *entFM* was not significantly different in BHI (p=0.7689), but was 5-fold higher in LB (p=0.0152). Mean RPKM values for the putative hemolysin A and the CDC cereolysin O genes were 58 and 103, respectively, in vitreous (Fig. 3d). The *hlyA* gene was 4-fold higher (p=0.0272), but *cerO* was not significantly different in BHI (p=0.8245) relative to the vitreous environment. In LB, *hlyA* was approximately 8-fold higher (p=0.0129), and while *cerO* trended towards higher levels of expression, this difference was not significant (p=0.1480) relative to vitreous.

**Figure 3.**
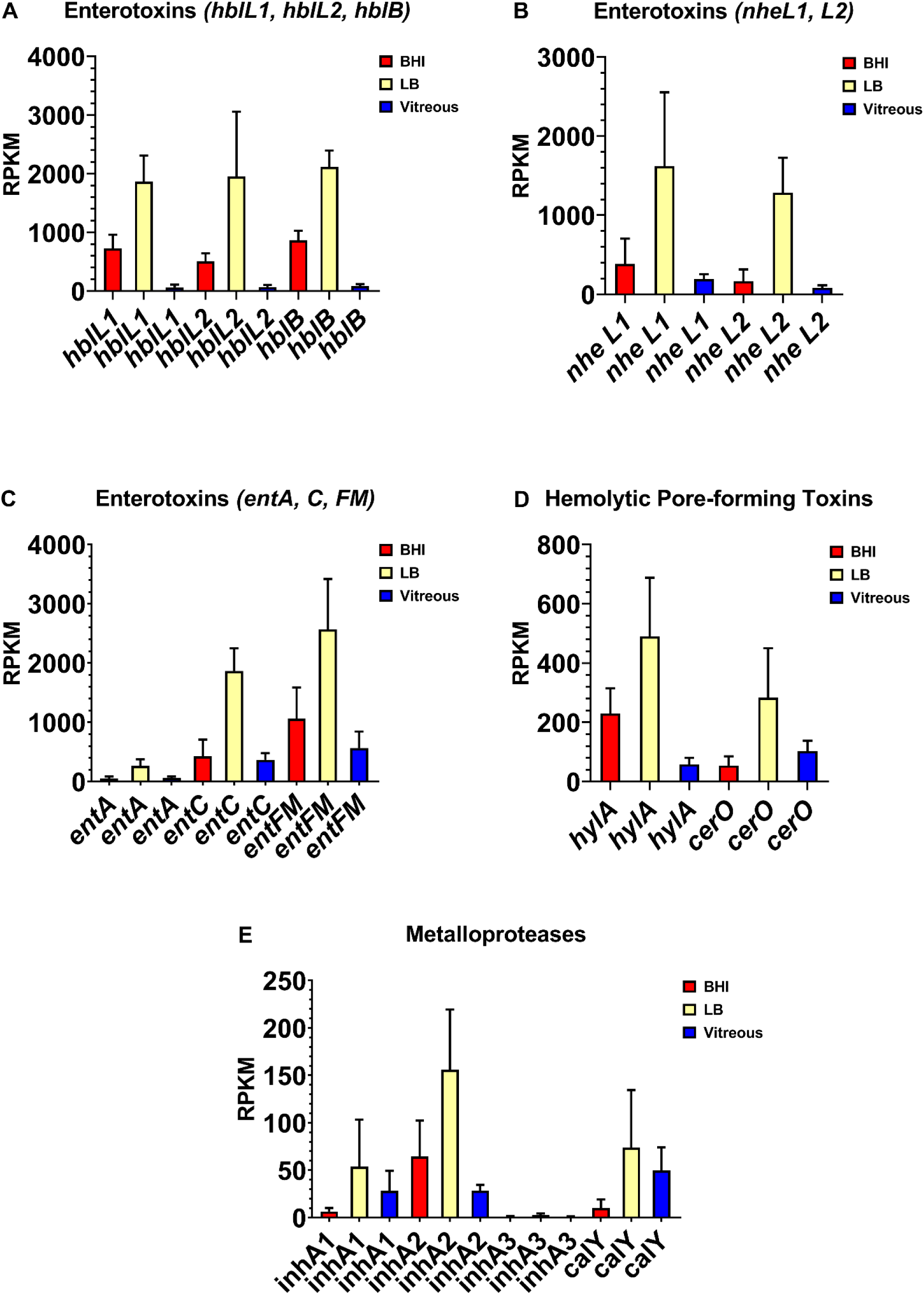
Normalized virulence-related gene expression in each of the three different environments. Reads per kilobase per million (RPKM) for the genes specifying hemolysin BL (Hbl) (A), nonhemolytic enterotoxin (Nhe) (B), enterotoxin FM and the putative enterotoxins A and C (C), the putative hemolysin A and the hemolysin cereolysin O (D), and the metalloproteases InhA1, InhA2, InhA3, and CalY (E). Brain Heart Infusion broth (BHI) is shown in red, Luria-Bertani broth in yellow, and vitreous in blue. RPKM values are the means ± the standard deviations of three independent RNA-Seq runs.

We also evaluated the expression levels of genes encoding the metalloproteases InhA1, InhA2, InhA3, and camelysin in the vitreous environment. The *inhA1* and *inhA2* genes were expressed in vitreous to a similar degree, each with a mean RPKM of 28 (Fig. 3e). There were no significant differences in *inhA1* expression between BHI, LB, or vitreous (p=0.2540). Transcript levels of *inhA2* in BHI was not significantly different from vitreous (p=0.5861), however was 6- fold higher in LB relative to vitreous (p=0.0250). In all three environments, expression of *inhA3* was not detected (Fig. 3e). The *calY* gene, which encodes a cell surface-associated metalloproteinase, camelysin, was expressed at similar levels to *inhA1* and *inhA2* in the vitreous with a mean RPKM value of 50 (p=0.3402) (Fig. 3e). Expression levels of *calY* were similar in BHI, LB, and vitreous (p=0.1972). These results indicate that with exception of *inhA3*, expression of both known and putative *B. cereus* virulence factors occurred in the vitreous environment, with expression levels being similar or lower than in BHI or LB.

### Virulence-related transcriptional regulatory gene expression in *ex vivo* vitreous

Genes reported to be involved in virulence factor regulation were expressed in vitreous during stationary phase, including the *sinR/sinI* system*, entD, codY, gntR, nprR*, and the *plcR/papR* system. Among these regulatory systems, the biofilm formation- and enterotoxin-related regulatory gene *sinR* was the most highly expressed regulator in the vitreous environment, with a mean RPKM value of 1164 (Fig. 4a). Interestingly, *sinR* expression was 7-fold higher in vitreous than in BHI (p= 0.0035), but not significantly different from expression in LB (p= 0.7855). Low level expression of the SinR inhibitor *sinI* was detected in the vitreous and BHI, with a mean RPKM value of 13 in both environments. *sinI* expression was undetected in LB (Fig. 4a). Mean RPKMs for *entD*, *codY*, *gntR*, and *nprR* were 25, 259, 193, and 44, respectively (Fig. 4b and c). In BHI, *entD*, *codY*, *gntR*, and *nprR* expression was not significantly different from vitreous (p≥0.0863). Expression of *entD* and *gntR* in vitreous and LB were similar (p≥0.1174). However, *codY* and *nprR* were 3-and 4-fold higher in LB relative to vitreous, respectively (p≤0.0394). Low level expression of the master toxin and virulence gene regulatory system *plcR/papR* was also detected in the vitreous. Mean RPKM values of 13 for *plcR* and 41 for *papR* were observed (Fig. 4d). Expression levels of these genes were not significantly different from both BHI (p≥0.2868) and LB (p≥0.2868). These results demonstrate that low, steady state levels of transcripts for virulence-related regulatory systems were detected in the vitreous during stationary phase, with most regulatory genes being similarly expressed in BHI, or lower than in LB. An important exception was *sinR*, which was significantly greater in both vitreous and LB relative to BHI. These results might suggest a role for SinR in a more nutrient limited environment of vitreous relative to nutrient rich BHI.

**Figure 4.**
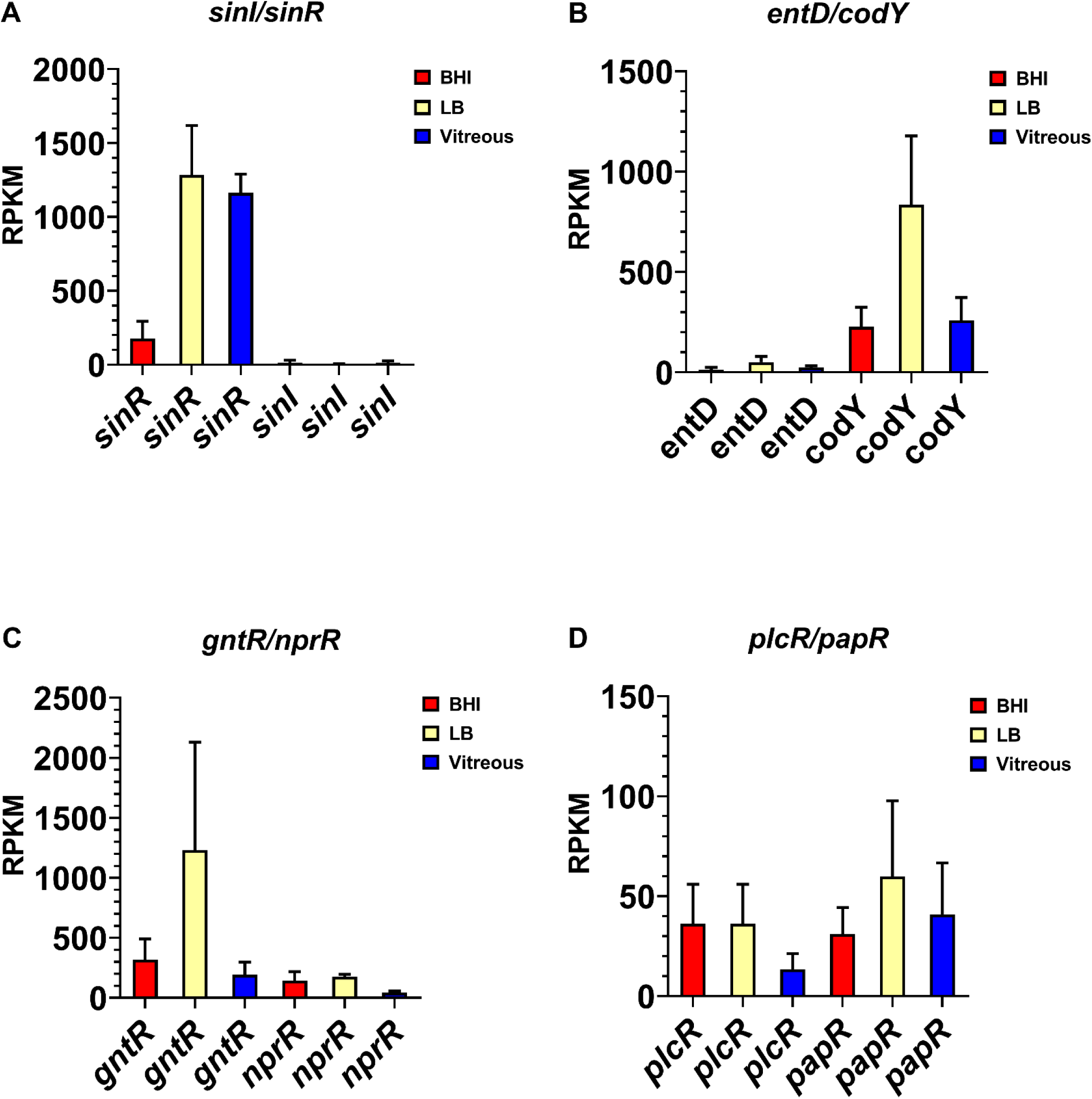
Normalized transcriptional regulatory gene expression in each of the three different environments. Reads per kilobase per million (RPKM) for the genes encoding transcriptional regulators SinI/SinR (A), EntD and CodY (B), GntR and NprR (C), and PlcR/PapR (D). Brain Heart Infusion broth (BHI) is shown in red, Luria-Bertani broth in yellow, and vitreous in blue. RPKM values are the means ± the standard deviations of three independent experiments RNA-Seq runs.

### Expression of genes associated with motility and chemotaxis in *ex vivo* vitreous

We reported that motility was an important phenotype for *B. cereus* in the eye, and that flagellar activity was essential for this process. However, in *ex vivo* vitreous, flagellar and chemotaxis gene expression was low to undetectable. Expression of the flagellar stator protein gene, *motB*, was not detected, and the mean RPKM values for the flagellar M-ring gene *fliF* and the flagellin subunit gene *fla* were 7 and 4, respectively (Fig. 5a). Expression of *fliF* was 7-fold higher in BHI relative to vitreous (p=0.0340), and 12-fold higher in LB relative to vitreous (p=0.0014). The expression of *fla* was not significantly different among the 3 environments (p=0.3034). The mean RPKM of *cheA*, *cheY*, and *cheR* was 5, 8, and 5, respectively, in vitreous (Fig. 5b).

**Figure 5.**
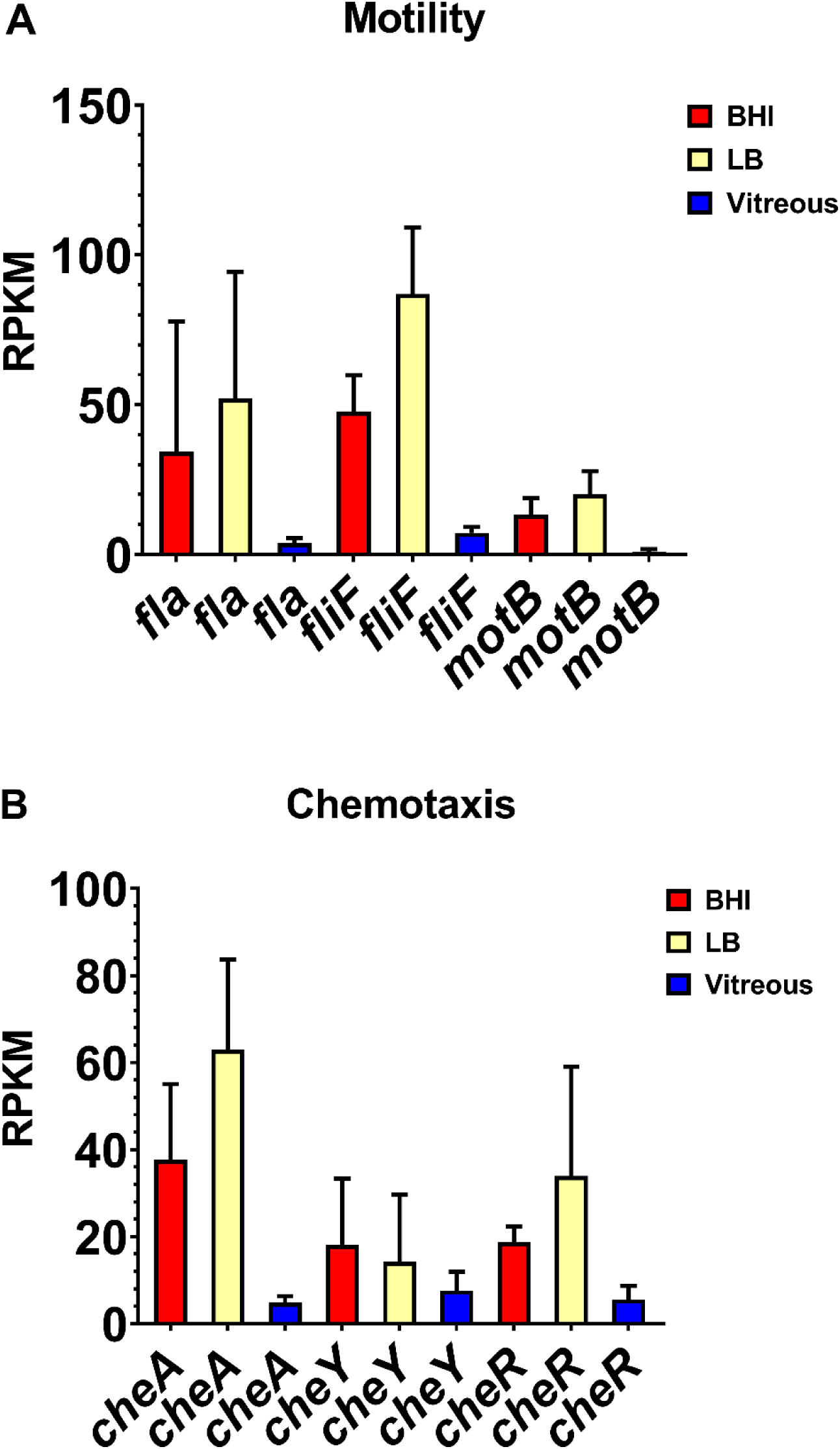
Normalized motility and chemotaxis-related gene expression in each of the three different environments. Reads per kilobase per million (RPKM) for motility-related genes *fla*, *fliF*, and *motB* (A), and chemotaxis-related genes *cheA*, *cheR*, and *cheY* (B). Brain Heart Infusion broth (BHI) is shown in red, Luria-Bertani broth in yellow, and vitreous in blue. RPKM values are the means ± the standard deviations of three independent RNA-Seq runs.

Expression levels of *cheA*, *cheY*, and *cheR* were not significantly different in BHI (p≥0.0922). In LB, expression of *cheA* was 13-fold higher relative to vitreous (p=0.0092), but neither *cheY* nor *cheR* expression was significantly different from vitreous (p≥ 0.1234). The lack of significant expression of chemotaxis and motility genes in *ex vivo* vitreous suggests that the *in vivo* environment may be necessary for higher level expression, as these genes are involved in the motility and migration of *B. cereus* within the eye during endophthalmitis.

### Expression of superoxide dismutase in *ex vivo* vitreous

Surprisingly, we discovered that the expression levels of two variants of the manganese superoxide dismutase gene, *sodA1* and *sodA2*, were among the most highly expressed genes during stationary phase in the vitreous compared to BHI and LB. The mean RPKM for *sodA1* was 599 and for *sodA2* was 3,326 after growth in vitreous (Fig. 6), the latter being the most highly expressed gene in the vitreous environment. Expression of *sodA1* and *sodA2* was not significantly different from vitreous after growth in BHI (p≥ 0.4278), or LB (p≥0.2936). The high level expression of *sodA2* in the vitreous suggested that superoxide dismutase production might be important during growth in the eye in order to protect *B. cereus* from internally-produced superoxide, or possibly from superoxide generated by neutrophils. In *S. aureus*, expression of the manganese superoxide dismutase gene *sodA* has been shown to be significantly upregulated after cultures were exposed to an internal superoxide generating agent [24]. Moreover, the viability a *sodA*-deficient *S. aureus* strain in stationary phase was 1000-fold lower than the parental wild type strain, and was significantly less virulent than the wild type strain in a mouse model of subcutaneous infection [24].

**Figure 6.**
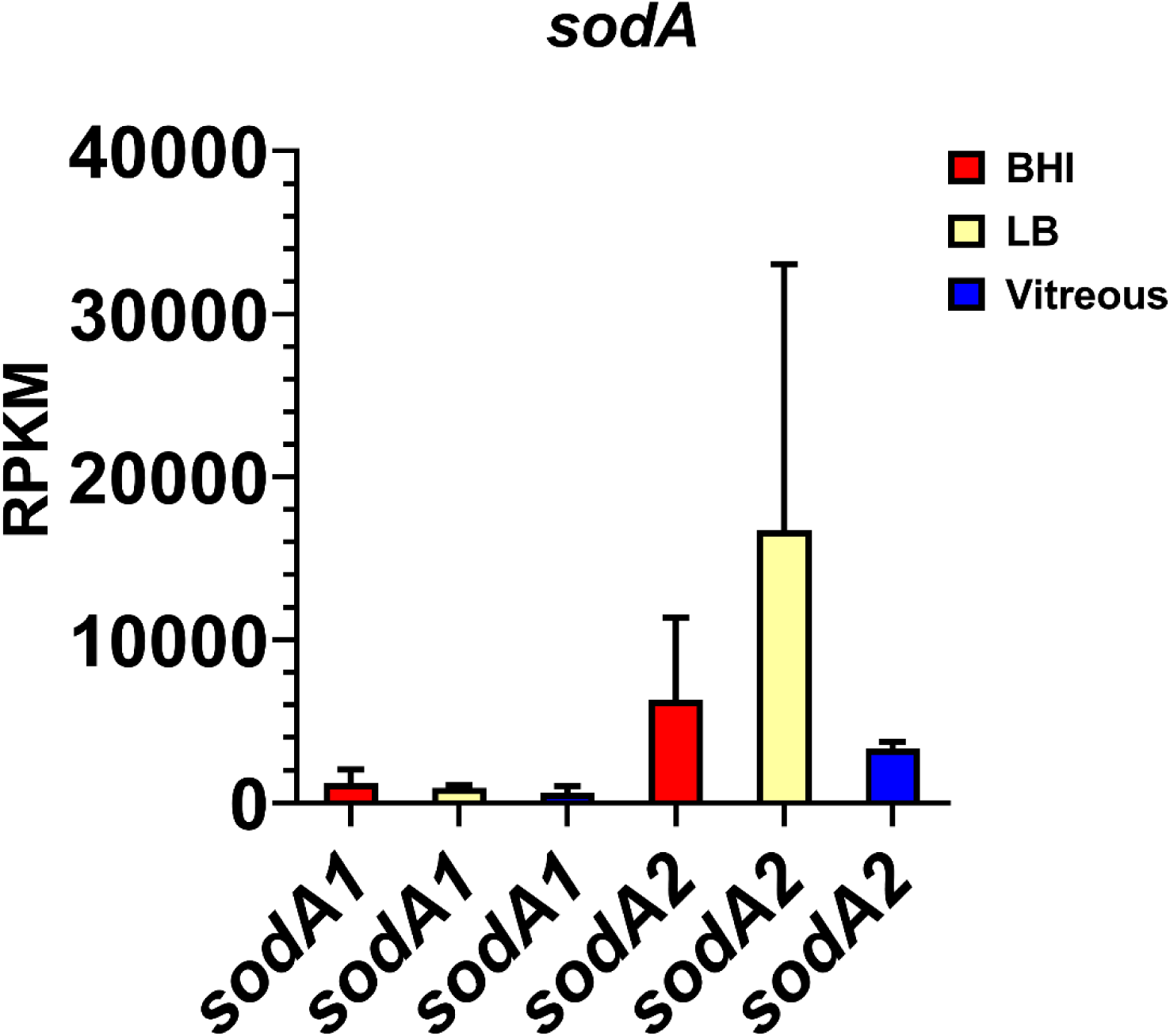
Normalized superoxide dismutase gene expression in each of the three different environments. Reads per kilobase per million (RPKM) for the genes *sodA1* and *sodA2*. Brain Heart Infusion broth (BHI) is shown in red, Luria-Bertani broth in yellow, and vitreous in blue. RPKM values are the means ± the standard deviations of three independent RNA-Seq runs.

## Discussion

To better understand the possible contributions of known and putative *B. cereus* virulence factors to endophthalmitis, we assessed the expression levels of a set of virulence factors, virulence-associated transcriptional regulators, and genes related to motility and chemotaxis after growth to stationary phase in vitreous compared to standard laboratory media. This time point was selected as it would correlate to later stages of infection when there is significant inflammation and damage to the retina. Both rabbit and mouse models of experimental *B. cereus* endophthalmitis have been used to elucidate the progression of intraocular infection. Bacteria replicate rapidly upon entry, and migrate throughout the posterior and anterior segments [12, 17, 23]. This rapid replication rate is likely attributable in part to the absence of immune responder cells. Inflammatory cells begin to enter the eye as early as 4 hours post-infection [23], however *B. cereus* replication remains unchecked. By 12 to 18 hours post-infection, the retinal architecture is completely disrupted and retinal function lost [23]. Cases of human *B. cereus* endophthalmitis also evolve rapidly and result in blindness or loss of the eye within 12 to 48 hours [2–5]. Evaluation of the expression levels of virulence factors in *ex vivo* vitreous late in the bacterial growth phase might identify factors that play key roles in interfering with immune cell function and suggest a possible mechanism for unchecked growth and lack of clearance of *B. cereus* from the eye.

The non-hemolytic enterotoxin (Nhe) and hemolysin BL are both pore-forming toxins that contribute to the pathogenesis of *B. cereus* gastroenteritis. While expression of both was detected in vitreous, levels were lower than in laboratory media. Hemolysin BL (Hbl) is a secreted dermonecrotic tripartite hemolysin consisting of two lytic components, L1 (*hblD*) and L2 (*hblC*), and a single binding component, B (*hblA*), located adjacent to one another on the genome [25, 26]. Hbl expression was observed in 42 to 73% of isolates that caused food poisoning [27, 28], and the *hblCA* genes were detected in approximately 15% of endophthalmitis and 37% of keratitis isolates [29]. Beecher et al. reported that purified Hbl injected into rabbit eyes elicited pathological effects clinically similar to those observed in *B. cereus* endophthalmitis, suggesting a possible role for Hbl [26]. However, in a rabbit model of *B. cereus* endophthalmitis, an overall similar course of infection was observed in rabbit eyes infected with either a wild type *B. cereus* or an isogenic Hbl-deficient mutant [11], suggesting that Hbl does not play an essential role in experimental *B. cereus* endophthalmitis [11]. The observation that outcomes were poor in animals infected with the Hbl-deficient *B. cereus* could relate to the production of additional toxins and the intense immense innate immune response that is not mitigated by the absence of Hbl. Here, Hbl was expressed in *ex vivo* vitreous; however, its cytotoxic effects might be masked *in vivo* by the plethora of other toxins produced. Non-hemolytic enterotoxin (Nhe), a tripartite enterotoxin similar to hemolysin BL, is also regulated by the PlcR/PapR system [30]. Nhe also requires the combined action of the three proteins, the lytic components NheA (*nheL2*) and NheB (*nheL1*), and the binding component NheC. In contrast to Hbl, Nhe is expressed in nearly all (97-99%) food poisoning-associated isolates [27, 28]. The role of Nhe has not been assessed in endophthalmitis, although, like Hbl, any effects on retinal or inflammatory cells might be masked by the activities of the other toxins produced in the eye.

Among the enterotoxins surveyed in this study, the *entFM* gene was the most highly expressed in the vitreous. EntFM was originally deemed an enterotoxin because of its ability to elicit fluid accumulation in rabbit and mouse ligated intestinal loops [31, 32, 33]. The *entFM* gene is found in both *B. cereus* and *B. thuringiensis*, and is associated with food-borne outbreak isolates [34, 35]. EntFM has been suggested to be a cell wall peptidase, and was demonstrated to induce vacuolization of macrophages, contribute to adhesion to HeLa cells and biofilm formation, and contribute to significant lethality in a *Galleria mellonella* model [36]. Regulation of *entFM* expression is independent of PlcR [37] which supports our finding of much higher transcript levels than the Hbl- and Nhe-associated transcripts in the vitreous. We also evaluated expression of two putative enterotoxin/cell-wall binding protein genes, *entA* and *entC*. While *entA* transcript levels were similar to that of the hbl-associated genes, *entC* was detected at 6-fold higher levels than *entA* in the vitreous. The role of *entFM*, *entA*, and *entC* in endophthalmitis has not been investigated.

Cereolysin O (CerO) is a pore-forming, heat-labile protein that is a member of the CDC family. While CerO requires cholesterol, there is no requirement for other specific cell-surface receptors, and CerO can lyse nearly all mammalian cells. CerO expression is controlled by the PlcR/PapR system [37]. However, the role of cereolyin O in the pathogenesis of *B. cereus* endophthalmitis remains unknown. We reported that Toll-like receptor 4 (TLR4) contributes to the robust inflammatory response to *B. cereus* during endophthalmitis, and identified a cohort of TLR4-dependent inflammatory mediators that are upregulated in the retina 4 hours following *B. cereus* infection [22]. This suggests that *B. cereus* is capable of directly or indirectly activating TLR4. Among its many ligands, TLR4 recognizes members of the CDC class of bacterial pore-forming toxins [38–41]. While CerO, a member of the CDC class, has not been shown to interact with and/or activate TLR4, our laboratory found that TLR4-dependent inflammatory mediators were significantly downregulated 4 hours following intraocular infection with a cereolysin O-deficient *B. cereus* strain [42]. In addition to direct cytotoxicity against cells in the retina, CerO might contribute to the rapid and destructive course of *B. cereus* endophthalmitis by activation of TLR4.

*B. cereus* secretes a battery of metalloproteases that that have been postulated to contribute to virulence by interfering with host defenses and degrading extracellular matrix components [43]. The *B. cereus* genome encodes three immune inhibitor A metalloproteases. InhA1 and InhA2 share 66% identity, InhA1 and InhA3 share 74% identity, and InhA2 and InhA3 share 71% identity [44]. All three proteins possess conserved zinc-binding and catalytic sites common to other known metalloproteases, and have cleavage sites, suggesting they are all secreted. Guillemet et al. observed low levels of expression of all three *inhA* genes in logarithmic phase and increased expression during stationary phase [44]. In our current analysis, we detected expression of *inhA1* and *inhA2* during stationary phase in the vitreous, but did not detect *inhA3* expression under these conditions. Guillemet et al. reported that a mutant strain of *B. cereus* deficient in all three *inhA* genes was attenuated in a *G. mellonella* insect model [44]. InhA hydrolyzes and inactivates cecropin and attacin, antibacterial proteins found in insect hemolymph [45]. *B. anthracis* InhA1 (91% identity with *B. cereus*) digests extracellular matrix proteins, fibronectin, laminins, and collagen I and IV [46]. This activity has been postulated to contribute to the ability of the bacteria to cross host barriers and invade deeper tissues [43]. Another putative role for these metalloproteases is in spore survival and escape from macrophages. Deletion of all three *inhA* genes did not reduce the cytotoxicity of *B. cereus* supernatant for macrophages *in vitro*. However, spores of the *inhA1* mutant, but not the *inhA2* or *inhA3* mutants, were incapable of escaping macrophages [44]. Therefore, InhA1 is apparently essential for efficient spore release from macrophages. In the eye, the InhA family of metalloproteases might collectively contribute to pathogenesis by multiple mechanisms. The InhAs might cause alterations in the vitreous by breaking down type II collagen, the predominant form of collagen comprising the vitreous [47]. Given that InhA1 facilitates the escape from macrophages, InhA1, and possibly InhA2 and InhA3, could function in a similar manner and promote escape from neutrophils. The casein-cleaving metalloproteinase, camelysin (*calY*), is cell-surface bound and has proteolytic activity towards serum protease inhibitors, collagen type I, fibrin, fibrinogen, and plasminogen [48, 49]. Transcript levels of the *calY* gene were similar to *inhA* transcript levels after growth in the vitreous environment. This raises the possibility that camelysin, similar to the InhAs, might contribute to the virulence of *B. cereus in vivo* by contributing to the breakdown of the vitreous by degrading collagen, and of the blood retinal barrier by degrading tight junction proteins.

In our study, we observed expression of genes involved in virulence factor regulation, including *sinR/sinI, entD, plcR/papR*, *codY, gntR,* and *nprR*. SinR/SinI are phase regulators expressed by *B. subtilis* and pathogenic members of the *B. cereus sensu lato* group. SinR has been reported to be involved in biofilm formation in *B. subtilis* and *B. thuringensis,* and enterotoxin production. Fagerlund et al. reported that SinR, along with PlcR, regulate the constitutive expression of Hbl in *B. thuringensis* biofilms [50]. In our analysis, SinR was expressed in all three *in vitro* environments. Surprisingly, *sinR* is one of the few genes that was expressed at higher levels in vitreous than in BHI. Interestingly, SinI, which represses *sinR* transcription [51], was expressed at low levels in BHI and vitreous, but was not detected in LB. The expression of *sinR* in the vitreous may result in constitutive expression of the *hbl* genes.

SinR may also regulate the expression of lipopeptides reported to be required for survival in other systems [50]. Another regulator involved in biofilm formation, EntD, was expressed in all three environments, albeit at low levels. EntD was the first Ent family protein identified in the *B. cereus sensu lato* group. EntD regulates virulence genes and genes with virulence-associated functions such as those involved in cell metabolism, cell structure, antioxidation, motility, and toxin production [52]. EntD regulates a four gene operon that synthesizes rare glycans present on the spore surface important for receptor recognition [53]. EntD is also involved in the regulation of proteins associated with the bacterial surface and are important in motility and biofilm formation. Expression of *entD* in the vitreous would be expected due to its role in regulating the expression of a number of *B. cereus* virulence factors. Expression of these factors involved in survival might be necessary to survive the stressful environment of the ocular environment, especially in the face of an innate immune response.

*B. cereus* is most known as the etiological agent of gastrointestinal diseases and local and systemic infections. *B. cereus* expresses a large number of virulence factors and toxins that result in the pathogenic effects of these diseases, most/many of which are under the regulation of quorum sensing. The PlcR/PapR is a pleiotropic quorum sensing system that activates the expression of a subset of virulence factors at the onset of stationary phase, while the Spo0A-AbrB regulatory circuit partially controls the plasmid-borne cereulide synthetase (*ces*) operon. The *plcR*/*papR* genes were detected at low levels across all three environments. The *plcR* gene is typically expressed under low nutrient conditions or when *papR* senses increasing cell density [30, 37]. Lereclus et al. reported that transient transcription of the *plcR* gene occurs at the end of the exponential growth phase in LB [54]. Expression was detected in stationary phase but only at low levels [54]. In our current study, expression levels of *plcR* and *papR* were similarly low in stationary phase in LB, and to a similar degree in BHI and vitreous. We previously demonstrated the importance of PlcR to the virulence of *B. cereus* in endophthalmitis [9]. Inflammation and retinal function loss were delayed by 18 hours in mouse eyes infected with a *plcR*-deficient strain relative to the wild type parental strain [9]. This suggests that *plcR* is expressed during infection of the eye. If the expression pattern of *plcR in vivo* and *ex vivo* vitreous is similar to that observed by Lereclus and colleagues in LB, then higher level expression would be detected earlier in infection or during exponential phase in *ex vivo* vitreous.

CodY, a nutrient-responsive regulator of Gram-positive bacteria, has a profound effect on both the PlcR/PapR and Spo0A-AbrB regulatory systems, which have been hypothesized to operate independently of each other [55]. CodY is a global transcriptional regulator of metabolism and virulence in low-GC Gram-positive bacteria. In *B. subtilis*, CodY regulates the expression of a myriad of genes in response to the nutritional status of the cell, including those for peptide transporters, proteases, and amino acid catabolism. CodY mainly functions as a repressor, repressing metabolic pathways when nutrients are available in excess. Frenzel et al. reported that deletion of *codY* resulted in downregulation of virulence genes belonging to the PlcR regulon and a concomitant upregulation of the *ces* genes [55]. Furthermore, CodY binds to the promoter of the immune inhibitor metalloprotease InhA1, demonstrating that CodY directly links *B. cereus* metabolism to virulence. *In vivo* studies using a *G. mellonella* infection model showed that the *codY* mutant was substantially attenuated, highlighting the importance of CodY as a key regulator of pathogenicity [55]. However, Sadaka et al. found that deletion of *codY* increased the virulence of *S. aureus* in a mouse model of anterior chamber infection [56]. While deletion of *codY* did not influence growth in the eye, inflammatory scores were significantly higher and retinal function retention was significantly lower after infection with the *codY* mutant relative to the wildtype parental strain [56]. These results suggested that in *S. aureus*, CodY downregulates target virulence genes during anterior chamber infection. We found that *codY* was expressed in BHI, LB, and in the vitreous environment. Expression during stationary phase in vitreous suggests a role in the upregulation of *inhA1* and involvement in the regulation of the PlcR regulon and its associated virulence factors. Another role for CodY might be to upregulate the expression of secreted proteases necessary to degrade the extracellular matrix, allowing *B. cereus* to breach the physical defenses in the eye. The possibility arises that in new environments such as vitreous or the eye, CodY regulates the expression of genes involved in metabolism necessary to utilize available nutrient sources.

GntR, the gluconate transcriptional repressor, was also expressed in all three environments *in vitro*. GntR is the negative transcriptional repressor of the gluconate operon (*gntRKPZ*), which encodes the proteins for gluconate utilization. GntR represses mRNA synthesis by binding to the *gnt* operator; the binding is suppressed by gluconate or glucono-delta-lactone. The *gnt* operon is involved in the ability of *B. subtilis* to utilize gluconate as a carbon source [57]. Expression of *gntR* in the vitreous would repress the expression of the other three genes necessary to metabolize gluconate, presumably due to the low concentration of or lack of gluconate in these environments. Gluconate is the product of oxidation of glucose, and can be readily utilized by organisms via the Entner-Doudoroff pathway. Gluconate is readily available in the intestinal tract and diarrheal pathogens such as *Vibrio cholera* and *Escherichia coli* adapt to the intestinal environment by its utilization [58]. Whether gluconate is available for utilization by *B. cereus* during endophthalmitis remains to be determined.

Expression of *nprR* was detected in all environments, albeit at lower levels than *codY* and *gntR*. NprR is a quorum sensing transcriptional regulator that regulates PlcR [59] at the beginning of stationary phase. NprR then regulates the expression of a number of genes involved in cell survival, sporulation, and antibiotic resistance, including the metalloprotease *nprA* gene [59]. The *nprR* gene is repressed by CodY during log phase growth. Detection of *nprR* expression in LB, BHI, and vitreous was not surprising considering that RNA sequencing was performed at a time point when cultures were in stationary phase and cellular density was high.

We have previously implicated the motility of *B. cereus* in the pathogenesis of endophthalmitis [8, 18] and were interested in examining expression levels of components of chemotaxis and motility during growth in vitreous. Genes associated with synthesis of the flagella and motor complexes, as well as chemotaxis related genes are located in an approximate 45 kb region in the *B. cereus* ATCC14579 genome [60, 61]. In *B. subtilis*, flagella- and chemotaxis-associated genes form part of a single operon that is regulated by CodY [62, 63]. In both *B. subtilis* and *B. cereus*, motility- and chemotaxis-associated genes are repressed under various stress conditions, including high salinity and the presence of bile salts [60, 64, 65]. Here, we observed low-level expression of the chemotaxis-related genes *cheA*, *cheY*, and *cheR*, and low to undetectable levels of the flagella- and motility-related genes *fla*, *fliF*, and *motB*. This contrasts with overall higher levels of expression of these genes at stationary phase in BHI and LB. CheA, CheY, and CheR comprise the regulatory system responsible for the switch between tumbling and forward movement and adjustment to chemoattract gradients [60, 65–68]. In *ex vivo* vitreous, nutrient distribution is likely to be homogenous and therefore the regulatory genes necessary for switching between tumbling and forward motion are not needed, given that chemotaxis is unnecessary. Synthesis of flagellar machinery would also not be necessary as there is no need for movement. Alternatively, a signal necessary for high-level expression of the chemotaxis- and motility-associated genes might be absent from the *ex vivo* vitreous environment.

Among the genes surveyed in our study, the *sodA2* gene was the most highly expressed in vitreous. In fact, *sodA2* was the most highly expressed of all *B. cereus* genes in this environment. Superoxide dismutases (SODs) are ubiquitous enzymes found in all kingdoms due to their importance in providing protection from oxidation. There are 4 families of SODs based on the metal cofactor required for optimal enzymatic activity: copper/zinc, nickel, manganese, and iron [69]. SodA2 is a Mn-based superoxide dismutase, and this type is found in bacteria, chloroplasts, mitochondria, and the cytoplasm of eukaryotic cells. SOD catalyzes the dismutation of superoxide anion (O2^-^) and other oxygen radicals to molecular oxygen and hydrogen peroxide (H2O2). Peroxidases and catalases then convert H2O2 to H2O and O2. In inflammatory diseases, O2^-^ activates the endothelium and attracts neutrophils. Neutrophils cross the endothelium and produce reactive oxygen species (ROS) which can damage host cells and tissues [70]. In *B. subtilis*, SodA is important in protection from oxidative stress in growing and sporulating cells [71]. Comparisons of the structures of *B. subtilis* SodA and human SOD revealed that human SOD possesses one less helix in a helical domain, a larger turn between antiparallel beta strands, and three different residues at the intersubunit interface [72]. These structural differences suggest that therapeutics targeting the *B. cereus* SodA2 would not interfere with the human SOD. In the eye, *B. cereus* production of SOD may inhibit or inactivate neutrophils, making the infection more difficult to clear. Conversely, inhibiting SOD may result in bystander damage from excess neutrophil activity.

These results provide insight into expression of virulence-related genes that might enable *B.cereus* to more readily adapt to the vitreous environment and establish endophthalmitis. These gene products might therefore serve as targets for adjunctive therapeutic agents. SOD represents a potential targetable candidate as it might function to interfere with neutrophil activity and prevent efficient clearance of *B. cereus* from the eye. Genes that we previously have shown to contribute to *B. cereus* virulence in the eye were not expressed or expressed at low levels in vitreous suggesting that additional factors present in the eye and/or an immune response might influence expression of these genes. Future studies will aim to characterize the *in vivo* virulome of *B. cereus* and identify genes that can be targeted to mitigate virulence.

## Author Statements

### Authors and contributors

PSC, FCM, and MCC designed the study. PSC, FCM, MAE, CL, ALL, and MHM performed the experiments. PSC, FCM, MAE, and CL compiled and analyzed the data. PSC performed the statistical analyses on the data. PSC and FCM prepared the original draft of the manuscript. FCM and MCC reviewed and edited the manuscript. PSC and MCC managed the project. MCC acquired the funding for this project. All authors read and approved the final manuscript.

## Acknowledgements

We thank Roger Astley (Department of Ophthalmology, OUHSC) and Mark Dittmar (Dean McGee Eye Institute Animal Facility) for their invaluable technical assistance. We thank Jenny Gipson and Allison Gillaspy at the Laboratory for Molecular Biology and Cytometry Research at OUHSC for assistance and the use of the Core Facility which provided the RNA-Seq service.

This work was presented in part at the 2017 American Society for Microbiology Microbe meeting in New Orleans, LA.

